# Analysis of SARS-CoV-2 specific T-cell receptors in ImmuneCode reveals cross-reactivity to immunodominant Influenza M1 epitope

**DOI:** 10.1101/2020.06.20.160499

**Authors:** John-William Sidhom, Alexander S. Baras

## Abstract

Adaptive Biotechnologies and Microsoft have recently partnered to release ImmuneCode, a database containing SARS-CoV-2 specific T-cell receptors derived through MIRA, a T-cell receptor (TCR) sequencing based sequencing approach to identify antigen-specific TCRs. Herein, we query the extent of cross reactivity between these derived SARS-CoV-2 specific TCRs and other known antigens present in McPas-TCR, a manually curated catalogue of pathology-associated TCRs. We reveal cross reactivity between SARS-CoV-2 specific TCRs and the immunodominant Influenza GILGFVFTL M1 epitope, suggesting the importance of further work in characterizing the implications of prior Influenza exposure or co-exposure to the pathology of SARS-CoV-2 illness.

---

In December 2019, SARS-CoV-2 or SARS-CoV-2, a novel coronavirus was first reported in Wuhan, China, and was later named a pandemic by the WHO in March 2020. In moderate to severe disease, the immune system is thought to play an important role in clinical presentation and course of illness (Qin et al. 2020; Zheng et al. 2020; Li et al. 2020; Shi et al. 2020). Given the extent of the morbidity and mortality that have been caused by SARS-CoV-2, there have been massive efforts across academia and industry to expedite our knowledge of the disease to develop better therapies and vaccines. In this effort, Adaptive Biotechnologies and Microsoft have partnered together to release ImmuneCode, a database of T-cell receptor sequences with derived specificities to SARS-CoV-2 epitopes through the use of the MIRA assay (Multiplex Identification of T cell Receptor Antigen Specificity) (Klinger et al. 2015). The assay exposes T-cells to antigens that span a significant portion of the viral genome and identifies antigen-specific TCR through the use of Next Generation Sequencing (NGS) TCR sequencing.

In this analysis, we wanted to query whether these SARS-CoV-2 specific TCRs were known to respond to other antigens. To do so, we collected known TCR-epitope pairs from McPas-TCR, a manually curated catalogue of pathology-associated T Cell Receptor Sequences and looked for SARS-CoV-2 specific TCRs within this database and further isolated the SARS-CoV-2 epitopes with cross-reactive TCRs to SARS-CoV-2 and other known antigens (Tickotsky et al. 2017). In our analysis, we reveal that there is notable cross-reactivity to the Influenza derived epitope M1 GILGFVFTL and these cross-reactive TCRs were present in both SARS-CoV-2 exposed and non-exposed individuals.

## Results

We first examined the distribution of TCRs within the McPas database over the types of pathogens present in the database and cross-referenced the SARS-CoV-2 specific TCRs into the McPas database (Figure 1A), and we noted that there was a statistically significant enrichment (from 17.3% to 32.9%) of SARS-CoV-2 specific TCRs that had known specificity to the immunodominant M1 GILGFVFTL epitope We then examined the distribution of TCRs within the ImmuneCode database across the various open readings frames (orfs) and mapped the M1 specific TCRs within this database (Figure 1B). We further noted that there was a statistically significant enrichment (from 11.8% to 66.7%) of cross-reactive SARS-CoV-2 specific TCRs within the surface glycoprotein orf. To better characterize the SARS-CoV-2 epitopes these TCRs were responsive to, we visualized the number of unique TCRs per epitope in the ImmuneCode database and noted a high level of specificity to a stretch of overlapping epitopes within the surface glycoprotein orf (Figure 1C). We observed that these 16 highly homologous TCRs (visualized by seqlogo) mapped to a set of overlapping epitopes containing an *SNVT* motif, suggesting that this repertoire of T-cells is capable of recognizing both the M1 epitope and this SARS-CoV-2-derived epitope. Finally, we found that these cross-reactive TCRs were present in both individuals who were and were not exposed to SARS-CoV-2 (Figure 1D).

**Figure 1.**
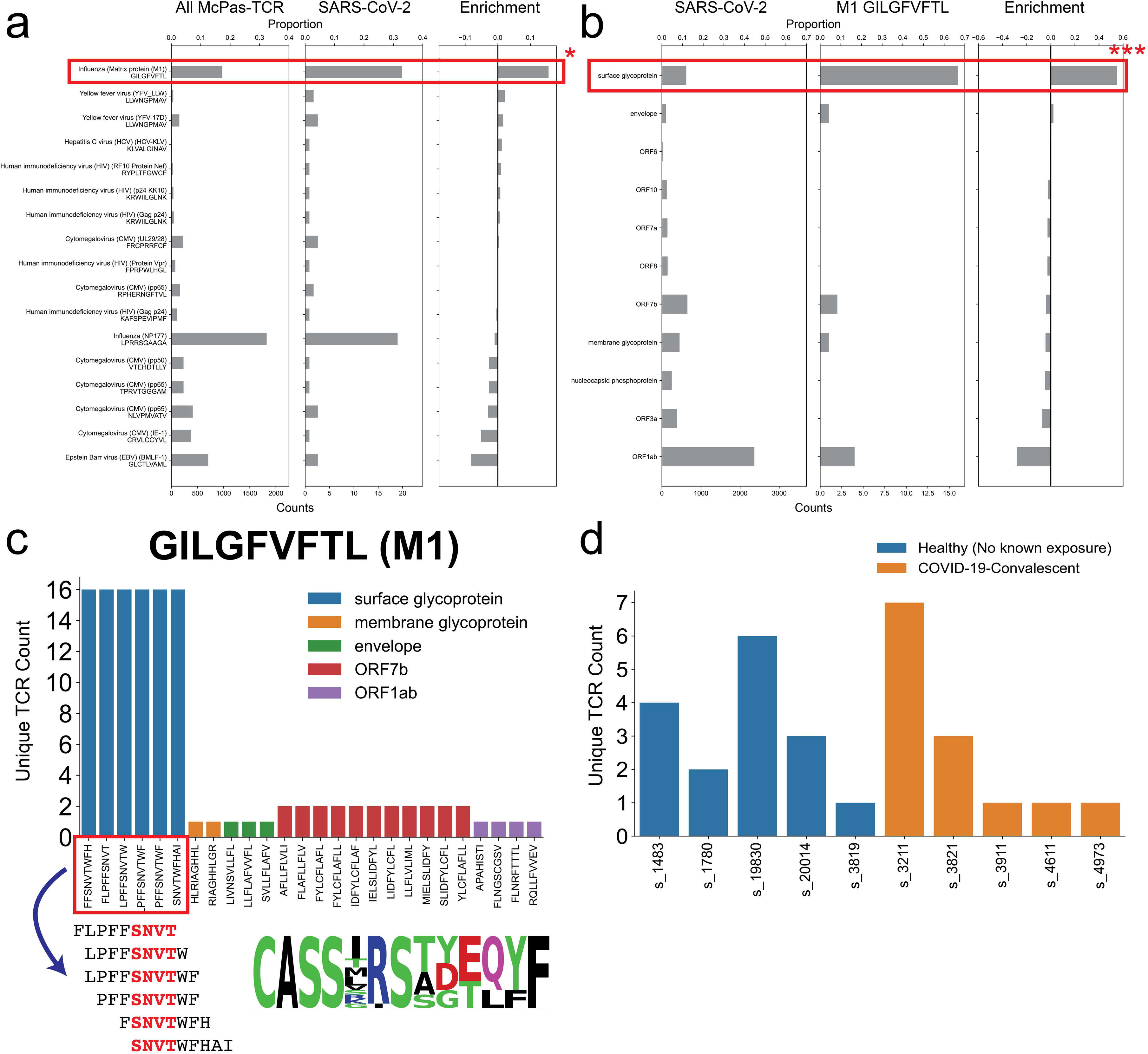
a) T-cell receptor sequences for pathogenic epitopes were collected from McPas-TCR and their distribution by pathogen was plotted (left) along with the distribution of SARS-CoV-2 specific TCRs (middle) that had exact sequence match to TCR sequences in the McPas-TCR database. Both counts and proportions for each pathogen is shown. The magnitude of enrichment is visualized (right) for each McPas-TCR epitope (Fisher’s Exact Test: *: p-val < 0.05 after multiple hypothesis testing correction) b) Distribution of SARS-CoV-2 TCRs from ImmuneCode (left) and M1 specific (middle) TCRs were visualized across all open reading frames (orfs). The magnitude of enrichment is visualized (right) for each orf (Fisher’s Exact Test: ***: p-val < 0.001 after multiple hypothesis testing correction). c) Distribution of 16 unique TCRs specific for M1 GILGFVFTL across SARS-CoV-2 epitopes (top). Select overlapping epitopes were highlighted showing a shared *SNVT* motif and corresponding specific TCRs were represented with a seqlogo (bottom). d) Distribution of 16 unique TCRs specific for M1 GILGFVFTL across individuals stratified by whether the individual was considered exposed or non-exposed to SARS-CoV-2.

## Discussion

In this brief analysis, we have identified a set of TCRs with known specificities to both a SARS-CoV-2 and Influenza antigen. The implications of these findings are relevant in understanding the nature of the immune response to SARS-CoV-2 as the clinical course is highly variable with patients presenting anywhere from being asymptomatic to requiring critical care (Goyal et al. 2020). Further insights into the evolution of the immune response could help guide early triage of individuals who are more likely to require more intensive support.

However, there are several limitations of this study. The number of individuals in this study was fairly small with 7 exposed and 5 non-exposed individuals. Given the lack of power in this analysis, it was difficult to assess for any differences in the extent of cross reactivity in exposed vs non-exposed individuals. Differences in cross reactivity may shed light on the role of Influenza specific antigen-experienced T-cells in the control of SARS-CoV-2 infection. Furthermore, given these results, future studies may want to consider comparing the clinical course in Influenza vaccinated vs unvaccinated individuals. Another significant limitation is assessing the certainty of antigen-specificity of a given TCR. Often, specificity of a TCR is determined in a high throughput fashion through methods such as tetramer sorting or T-cell expansion based assays, which can be inherently noisy and non-specific sources of data (Sidhom et al., n.d.). This problem is highlighted in Figure 1A when we compare the distribution of unique TCR sequences in the McPAS-TCR database vs the sampled distribution of SARS-CoV-2 specific TCRs. While there is clear enrichment in the M1 epitope, the NP 177 epitope, while demonstrating a large number of TCRs that are seemingly cross-reactive, the enrichment over the background distribution within McPas-TCR is not significant, suggesting that this may be nonspecific signal within the MIRA assay. In light of these limitations, T-cell recognition assays such as ELISPOT would be needed moving forward to validate these results, and confirm that truly, Influenza specific T-cells are capable also of recognizing SARS-CoV-2 specific epitopes. Finally, while these SARS-CoV-2 epitopes may elicit *in-vitro* responses, if they are not processed and presented *in-vivo*, Influenza specific T-cells would likely not have any ability to recognize SARS-CoV-2 infected cells. Thus, further work is required to not only confirm that these Influenza specific T-cells are cross-reactive to these SARS-CoV-2 epitopes but studies are needed to confirm their *in-vivo* processing and presentation.

In conclusion, while these results are preliminary in a small cohort of individuals, we have identified a set of TCRs that is known to both recognize an immunodominant epitope derived from Influenza and SARS-CoV-2, suggesting that immune control of one infection may play a role in the control of the other. These results justify further study into evolution of the immune response to SARS-CoV-2 in the setting of prior or coexisting Influenza infection.

## Data Availability & Code

All data and code used to conduct this analysis are available at https://github.com/sidhomj/COVID19 with included jupyter notebook to reproduce all analyses and figures present in this manuscript.

## References

Goyal, Parag, Justin J. Choi, Laura C. Pinheiro, Edward J. Schenck, Ruijun Chen, Assem Jabri, Michael J. Satlin, et al. 2020. “Clinical Characteristics of Covid-19 in New York City.” The New England Journal of Medicine 382 (24): 2372–74.

Klinger, Mark, Francois Pepin, Jen Wilkins, Thomas Asbury, Tobias Wittkop, Jianbiao Zheng, Martin Moorhead, and Malek Faham. 2015. “Multiplex Identification of Antigen-Specific T Cell Receptors Using a Combination of Immune Assays and Immune Receptor Sequencing.” PloS One 10 (10): e0141561.

Li, Xiaowei, Manman Geng, Yizhao Peng, Liesu Meng, and Shemin Lu. 2020. “Molecular Immune Pathogenesis and Diagnosis of COVID-19.” Journal of Pharmaceutical Analysis 10 (2): 102–8.

Qin, Chuan, Luoqi Zhou, Ziwei Hu, Shuoqi Zhang, Sheng Yang, Yu Tao, Cuihong Xie, et al. 2020. “Dysregulation of Immune Response in Patients with COVID-19 in Wuhan, China.” Clinical Infectious Diseases: An Official Publication of the Infectious Diseases Society of America, March. https://doi.org/10.1093/cid/ciaa248.

Shi, Yufang, Ying Wang, Changshun Shao, Jianan Huang, Jianhe Gan, Xiaoping Huang, Enrico Bucci, Mauro Piacentini, Giuseppe Ippolito, and Gerry Melino. 2020. “COVID-19 Infection: The Perspectives on Immune Responses.” Cell Death and Differentiation 27 (5): 1451–54.

Sidhom, John-William, H. Benjamin Larman, Petra Ross-MacDonald, Megan Wind-Rotolo, Drew M. Pardoll, and Alexander S. Baras. n.d. “DeepTCR: A Deep Learning Framework for Understanding T-Cell Receptor Sequence Signatures within Complex T-Cell Repertoires.” https://doi.org/10.1101/464107.

Tickotsky, Nili, Tal Sagiv, Jaime Prilusky, Eric Shifrut, and Nir Friedman. 2017. “McPAS-TCR: A Manually Curated Catalogue of Pathology-Associated T Cell Receptor Sequences.” Bioinformatics 33 (18): 2924–29.

Zheng, Meijuan, Yong Gao, Gang Wang, Guobin Song, Siyu Liu, Dandan Sun, Yuanhong Xu, and Zhigang Tian. 2020. “Functional Exhaustion of Antiviral Lymphocytes in COVID-19 Patients.” Cellular & Molecular Immunology.

